# Effects of Gold Nanoparticles on the Stepping Trajectories of Kinesin

**DOI:** 10.1101/2020.02.05.935791

**Authors:** Sabeeha Hasnain, Mauro L. Mugnai, D. Thirumalai

## Abstract

Substantial increase in the temporal resolution of the stepping of dimeric molecular motors is possible by tracking the position of a large gold nanoparticle (GNP) attached to a labeled site on one of the heads. This technique was used to measure the stepping trajectories of conventional kinesin (Kin1) using the time dependent position of the GNP as a proxy. The trajectories revealed that the detached head always passes to the right of the head that is tightly bound to the microtubule (MT) during a step. In interpreting the results of such experiments, it is implicitly assumed that the GNP does not significantly alter the diffusive motion of the detached head. We used coarse-grained simulations of a system consisting of the MT-Kin1 complex with and without attached GNP to investigate how the stepping trajectories are affected. The two significant findings are: (1) The GNP does not faithfully track the position of the stepping head. (2) The rightward bias is typically exaggerated by the GNP. Both these findings depend on the precise residue position to which the GNP is attached. Surprisingly, we predict that the stepping trajectories of kinesin are not significantly affected if, in addition to the GNP, a 1 *µ*m diameter cargo is attached to the coiled coil. Our simulations suggest the effects of the large probe have to be considered when inferring the stepping mechanisms using GNP tracking experiments.

**Statement of Significance:** Kinesin walks on microtubules by taking rapid steps separated by long dwells. The spatiotemporal resolution necessary to track the location of the motor during the fast displacements has been recently achieved by labeling one of the heads with a gold nanoparticle (GNP), and monitoring the light scattered by the probe. The results of the measurements depend on two crucial assumptions that currently cannot be verified with independent experiments: (i) the GNP faithfully tracks the motor head, and (ii) the giant probe does not alter the stepping trajectory. Using well-calibrated simulations, we show that the actual stepping trajectories of kinesin are not accurately reported by the probe, and hence, the inferences based on such experiments are not quantitatively accurate.

**Author Contribution:** SH, MLM, and DT designed the research. SH performed the simulations. SH, MLM, and DT analyzed the data and wrote the paper.

## Introduction

Conventional Kinesin (Kin1) is a processive motor protein that takes multiple, nearly uniform steps on the cytoskeletal track microtubule (MT) (1). Motors belonging to the kinesin superfamily are responsible for a number of cellular functions, such as cargo transport and chromosome separation during cell division (2–4). Kin1 moves towards the plus end of the MT (5–7) by hydrolyzing one ATP molecule per step (1, 5, 8–11). The two heads of the motor, joined through a coiled-coil tail domain, walk alternatively (the so-called “hand-over-hand” mechanism) by taking discrete ∼16.4 nm steps (1, 12, 13), resulting in a net average center of mass displacement of 8.2 nm. The step size is commensurate with the distance between two adjacent *αβ* tubulin dimers, which are the building blocks of the MT polymer (14–17).

Thanks to a remarkable series of experiments spanning over 25 year (7, 14, 18–23) many aspects of the stepping kinetics of kinesin are well established. In the absence of resistive force, the unidirectional stepping of Kin1 starts with the detachment of the trailing head (TH) from the MT. Binding of ATP to the leading head (LH) triggers neck linker (NL) docking to the cover strand, which is a central *β*-sheet of the LH (24–30). Docking of the NL minimizes the probability that the TH takes side steps on the neighboring MT protofilaments (31). A step is completed after (i) hydrolysis of ATP at the bound head (which may be coupled to the completion of NL docking (32)), (ii) ADP release from the unbound head, and (iii) reattachment of the TH to the target binding site (TBS), located ≈ 16.4 nm ahead of the initial binding site (32–34).

Although the global events in the unidirectional motion of Kin1, summarized above, are well-accepted, the precise mechanism of the stepping process continues to be a topic of ongoing debate. For instance, how kinesin waits to bind is controversial (34, 35). Several experiments have suggested that at high ATP concentrations, the detachment of the TH occurs only after ATP binds to the bound LH (1, 7, 12). In other words, ATP binding occurs in the two-head bound (2HB) state, when both the TH and LH are attached to the MT. In contrast, other studies, performed at low or limiting ATP concentrations, suggest that ATP binds only after TH detaches from the MT, that is in the one-head bound (1HB) state (23, 35, 36).

It is important to understand how the experimental techniques used to probe the stepping mechanism of Kin1 affect the trajectories explored by the TH as it searches for the TBS. Experiments using optical trapping assays, which monitor the movement of a cargo attached to the coiled-coil (14, 19, 20, 22, 37, 38), failed to detect the 1HB state possibly because of limited temporal resolution. Single-molecule fluorescence resonance energy transfer experiments have shown that, at low (≈ 10 *µ*M) ATP (≈ 10 *µ*M) concentrations, ATP binding to the LH is so slow that the motor waits in the 1HB state, which necessarily would require spontaneous detachment of the TH (31, 35).

Two recent insightful experiments, using similar protocols, have produced stepping trajectories of Kin1 as a function of ATP concentration at higher temporal resolution than previously possible. Mickolajczyk *et al*. (34) used interferometric scattering microscopy (iSCAT) (39–42) to track the motion of a 30 nm diameter gold nanoparticle (GNP) attached to one of the motor heads. They suggested that the TH detaches only after ATP first binds to the LH for ATP concentrations larger than 10 *µ*M. In contrast, Isojima *et al*. (35) used total internal reflection-based dark-field microscopy (DFM) to track the motion of a 40 nm GNP at nanometer precision and temporal resolution of the order of ≈ 55 *µ*s. The stepping trajectories recorded in this experiment were used to infer that ATP binds to the LH only after the TH detaches from the MT, that is in the 1HB state (see figure 4a in ref. (35)). Strikingly, the experiments performed by Isojima *et al.* (35) indicated that the detached head *always* passes the bound head to the right, which implies that the transverse coordinate (Y-axis in figure 1) of the TH is predominantly negative. Given the possibility of arriving at different conclusions from similar experiments, it is important to take into account measurement processes in analyzing the stepping trajectories not only in kinesin but also other motors.

**Figure 1:**
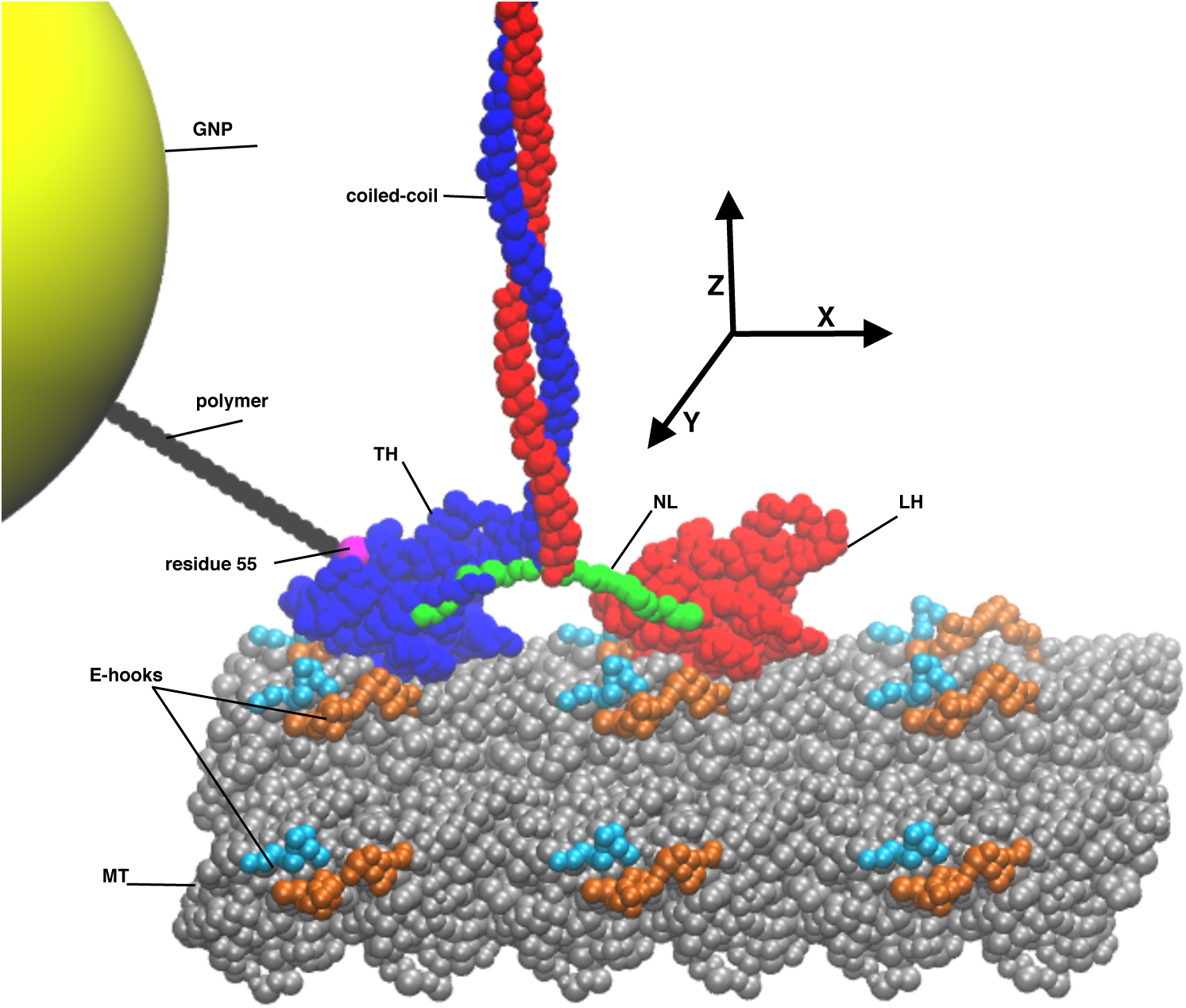
Sketch of the MT-Kinesin complex with GNP (shown in yellow) attached to residue 55 (shown in magenta) of the TH. A polymer (shown in black), consisting of 17 beads, connects residue 55 and the GNP. The neck linkers (NLs) of the TH and the LH are shown in green, and the charged E-hooks are in cyan and orange. The MT-axis is along the X-direction, the Y-direction corresponds to the lateral positions in the neighboring protofilaments, and the Z-axis is drawn perpendicular to the plane containing X and Y-axis. For clarity all the relevant parts of the complex are labeled.

**Figure 2:**
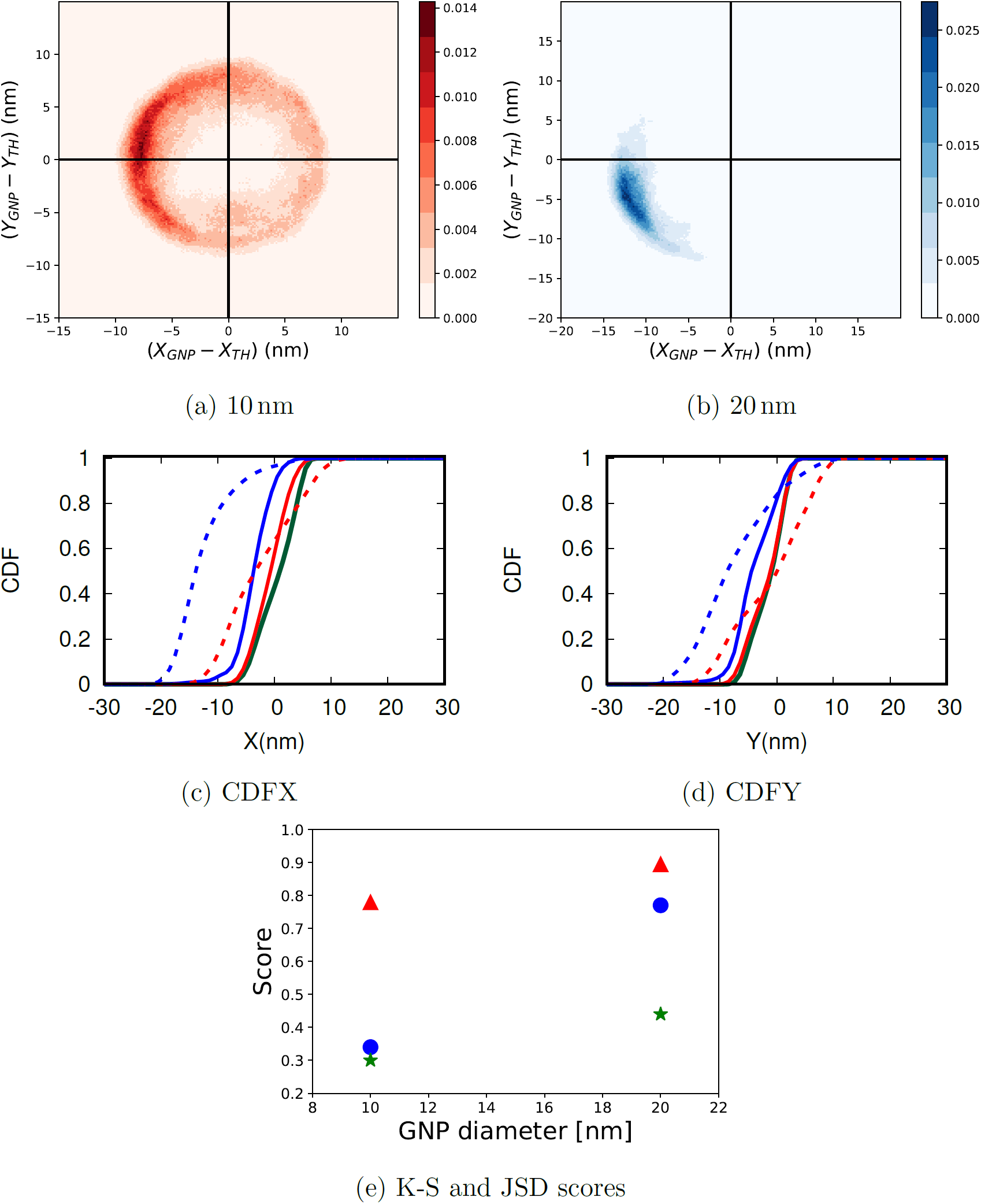
Distribution of the position of the GNP relative to the position of the TH projected onto the *X* and *Y* coordinates. (a) Density plot for a 10 nm-GNP, and (b) for a 20 nm-GNP. The scales on the right indicate the value of the probability density. (c) Cumulative Distribution Function (Eq.1) of the TH and GNP as a function of the *X* coordinate (parallel to the MT). The continuous lines refer to the TH, and the dashed lines correspond to the GNP. The green curve refers to the CDF for the TH in the absence of GNP, the CDF results for a 10 nm-GNP are reported in red, blue indicates simulations performed with a 20 nm-GNP. (d) Same as (c) except these are for the *Y* coordinate (transverse to the MT). (e) Results of the K-S (blue color is for the *X* direction, the *Y* direction is indicated by green stars). The results using the JSD (red triangles) measures (Eq.4) are given by the red triangles. Both the statistical tests show that GNP does distort the kinesin stepping trajectories.

**Figure 3:**
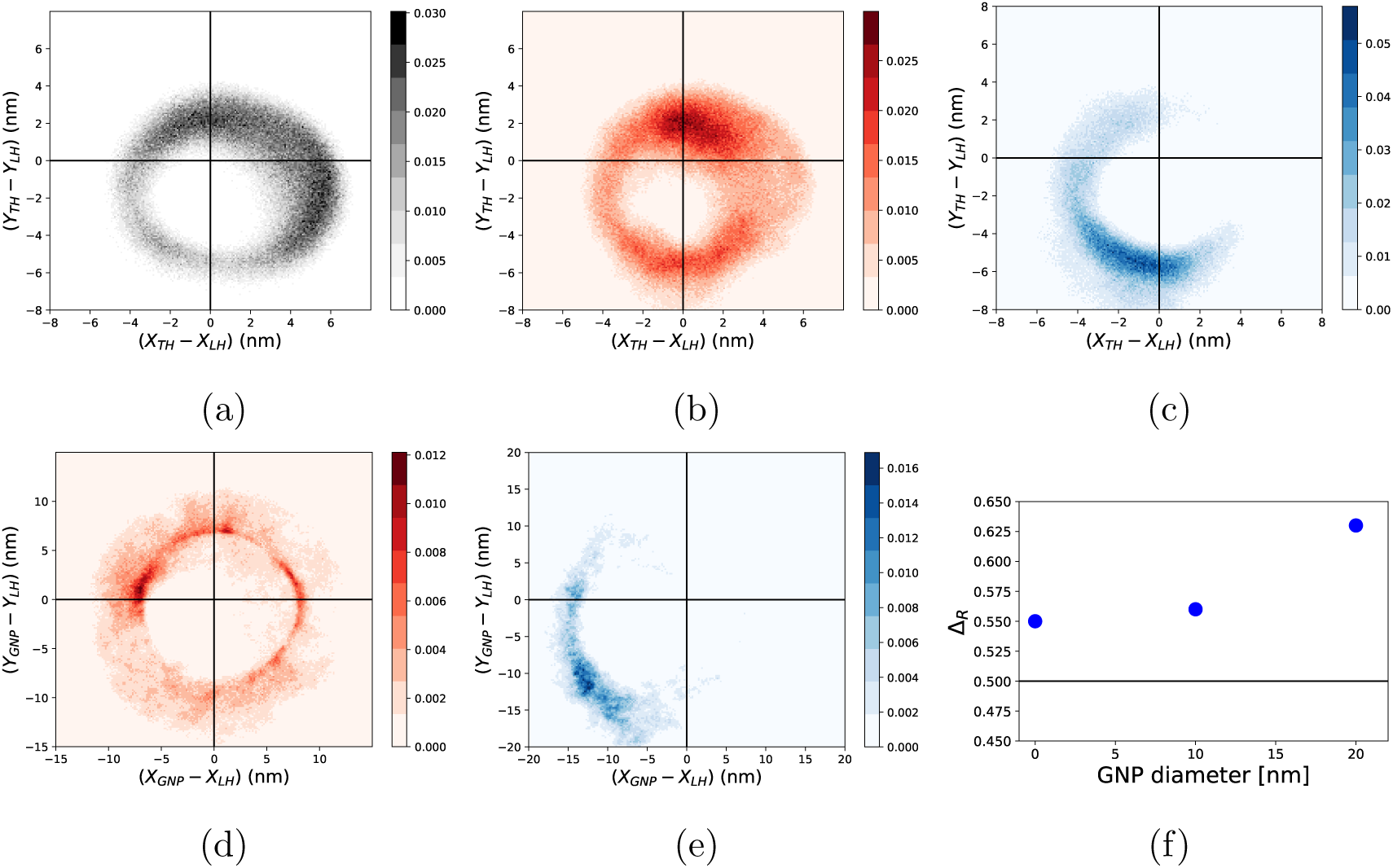
Two-dimensional distribution of the position of the TH of the motor or the GNP with respect to the LH. The black lines indicate *X =* 0 and *Y =* 0. Each panel shows a two-dimensional intensity plot, with the intensity scale given in the color bar on the right of each plot. (a)-(c) TH position relative to the (MT-bound) LH when there is no GNP (a), in the presence of a 10 nm GNP (b), and in the presence of 20 nm GNP (c). (d)-(e) GNP position with the respect to the (MT-bound) LH. The two panels (d) and (e) refer to simulations performed with (d) a 10 nm GNP, (e) a GNP of 20 nm. (f) Δ_*R*_ values for the TH without GNP, 10 nm and 20 nm GNP attached to residue 55. The black line correspond to Δ_*R*_ = 0.5.

**Figure 4:**
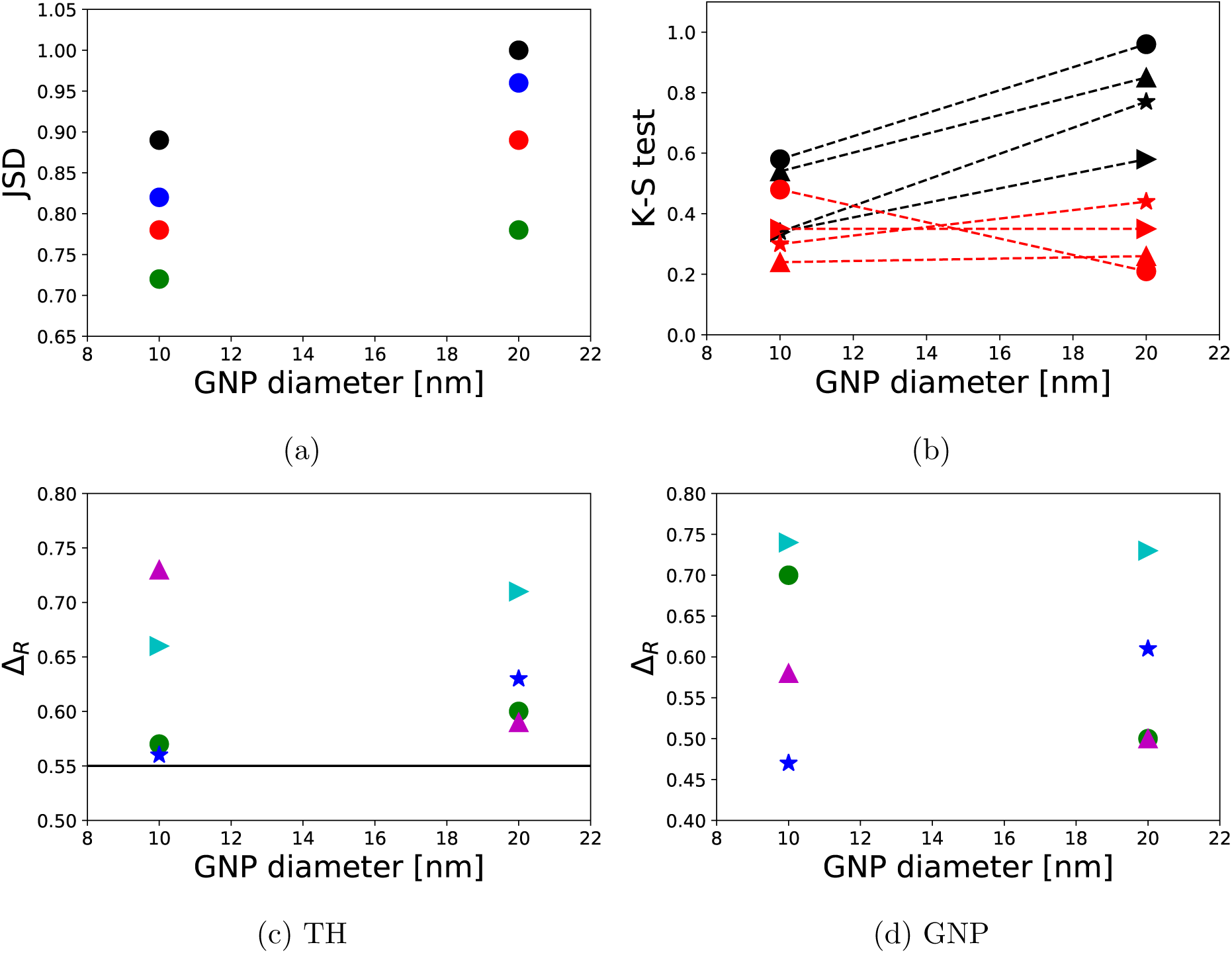
(a) The JSD scores for GNP attached to different residues on the TH. The points marked in black, blue, red and green represent the system where GNP is attached to residue 1, 55, 215, and 356, respectively. Note that the JSD score would be close to zero if GNP tracked the TH faithfully. (b) Same as except the results are the scores obtained from the K-S test. The points in black report the scores from the distribution along the X-direction, and the points marked in red show the score obtained from the distribution along the Y-direction. The circles (•) represent system with attachment of GNP at residue 1, stars (⋆) for attachment at residue 55, triangles (▴) for attachment at residue 215, and arrows (▸) for attachment at residue 356. Δ_*R*_ values for the position of the TH of the motor relative to the bound head. The black line corresponds to the system without GNP. The points represent system with attachment at residue 1 (•), 55 (⋆), 215 (▴), and 356 (▸). (d) The values of Δ_*R*_ calculated using the position of the GNP relative to the bound head. The points represent system with attachment at residue 1 (•), 55 (⋆), 215 (▴), and 356 (▸)

Here, we investigate the effects of attaching a GNP to one of the motor heads on the stepping trajectories by using a modification of a previously introduced coarse-grained (CG) model of the MT-Kin1 complex (31, 43). We address the following questions: (1) Does the experimentally measured trajectory generated by monitoring the position of the GNP faithfully reflect the location of the tethered head? An affirmative answer to this question is implicit in the analysis of the experimental data (34, 35). (2) Is the rightward bias in the movement of the TH found by Isojima *et al.* (35) a consequence of attaching a large GNP? (3) How does cargo affect the stepping trajectory of the TH when the GNP is attached? GNP assays conducted in the absence of cargo do not reflect the geometry of kinesin as a transporter. It is therefore interesting to examine the extent to which GNP tracking alters the real time trajectory of the stepping motor with cargo attached to the coiled coil (see figure 1). Because even in these high spatial and temporal resolution experiments the positions of the GNP and the TH cannot be simultaneously measured, computer simulations using models that are well calibrated against experiments (31, 43), might provide answers to these questions.

We found that (1) the movement of the GNP along the MT (*X*-axis in figure 1) does not fully track *X*_*TH*_, the projection of the TH coordinate onto the MT axis. Moreover, (2) the rightward biasing of the TH, which has been found to be the inherent property in the experiment that track the GNP, is enhanced in the presence of GNP. The extent of rightward bias in the TH of the motor during stepping is independent of the position of the attachment of the GNP, and increases with the size of the GNP. Finally, (3) we report a prominent role played by the cargo attachment to the coiled coil in dictating the trajectory of the tethered head even in the presence of the GNP. Surprisingly, we find that in the presence of cargo the extent of rightward bias with or without the GNP is similar. Experiments conducted on a cargo-loaded kin1 in the presence of the GNP could be used to test our predictions.

## Results

We performed a number of simulations in order to assess the impact of attaching the GNP to various residues of the TH on the kinesin stepping trajectories. The simulations were performed using the Self-Organized Polymer (SOP) coarse-grained model (44) for a system consisting of the MT-kin complex, coiled-coil, and the GNP attached to a specific residue of the TH (figure 1). The details of the energy function and the simulation methodology are provided in the Methods section. We performed the following set of simulations in order to address the question posed in the introduction.

A. We first generated stepping trajectories without the GNP, which serve as reference for evaluating the impact of the GNP giant probe on kinesin stepping. Our previous studies, which simulated MT-kin complex attached to the coiled coil but without the GNP (figure 1c of ref. (43)), showed that the simulations were in good agreement with experiments.
B. Next, we modeled the iSCAT and DFM experiments by attaching a GNP to the TH of the kinesin dimer. We repeated the simulations by varying the diameter of the GNP in the range 10 − 40 nm, which captures the sizes used in experiments. However, because the time for moving substantial distance along the MT axis for a 40 nm-GNP are much longer than the simulation times, our conclusions in this case are not quantitative.
C. In order to faithfully recapitulate the experimental setups, we repeated our simulations by attaching the GNP to different TH residues: 1, 55, 215, and 356.Attachment of the GNP was used in the experiments by Mickolajczyk *et al*. (34). In the Isohima *et al.* (35) experiments the GNP was attached to residues 55, 215, or 356. A typical simulation setup is portrayed in figure 1. By comparing simulations with and without GNP, the effects of GNP on stepping trajectories can be quantitatively assessed.
D. Finally, we tested the interplay between cargo and the GNP using a setup in which both of them are present. The results in this case are predictions, which will have to wait experimental validation.

Most of the results presented in the main text are obtained using the setup in which the GNP is attached to residue 55 of the TH. The analyses of the effects on kinesin motility resulting from attaching the probe to alternative residues are relegated mainly to the discussion section.

### Does the GNP faithfully trace the trajectory of the TH?

A major assumption in the interpretation of the experiments is that the paths traced by the GNP (almost) mirror the movement of the TH. In order to determine the extent to which this assumption is valid, we performed different tests to check if the GNP tracks the TH precisely. We calculated the distribution of the position of the GNP with respect to the location of the TH at each time step using 50 stepping trajectories of ≈ 25 *µ*s each. For a 10 nm GNP, attached to residue 55 of the TH, the GNP is predominantly localized on a circle of radius ≈ 7 nm around the TH (see figure 2a). When the size of the is increased to 20 nm, the diffusive motion of the GNP relative to the TH is confined to an arc of a ≈ 12 nm circle around the TH (figure 2b) on the time scale of our simulations. Therefore, from the distribution of the position of the GNP relative to the TH we cannot predict the exact position of the one with respect to the other.

#### Kolmogorov-Smirnov (K-S)Test

To quantify the difference in the distribution of GNP and TH we performed the K-S test (45) using the cumulative distributions of the position of the TH and GNP projected along the MT axis (*X* in figure 1) and relative to the location of the LH at *t* = 0. The cumulative distribution functions are given by,

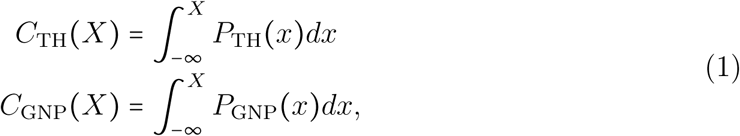

where *P*_*O*_ (*x*) refers to the probability density of finding *O* (*either*TH or GNP) in an interval *dx* around *x*. In the K-S test, the distance between the two CDFs is given by *d*_*X*_,

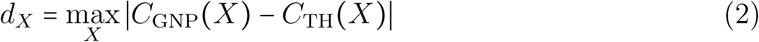

where *d*_*X*_ = 0 means that the two distributions are identical, whereas *d*_*X*_ = 1 indicate that there is no superposition between the two.

The CDFs shown in figure 2c, and the result of the K-S test in figure 2d illustrate that the GNP does not faithfully track the TH. In order to account for the transverse coordinate, we repeated the K-S test using the *y* coordinate to compute the CDF. Again, the location of both the TH and the GNP are computed in a frame of reference centered around the location of the MT-bound LH at *t* = 0. In this case, the K-S distance is given by,

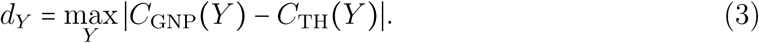

Just as shown in figure 2c the results in figure 2d indicate that there are differences between the CDFs computed in the absence and in the presence of GNP, which is clearly revealed in the K-S measure (figure 2e). Furthermore, the K-S scores (figure 2e) increase as the size of the GNP increases, indicating that larger GNP will track the position of the TH even less faithfully.

#### Jensen-Shannon Divergence (JSD)

To further characterize the ability of the GNP to monitor the location of the TH, we calculated the probability distribution function of the location of the TH and GNP along the X- and Y-directions relative to the position of the MT-bound LH. To measure the similarity between these two distributions, we calculated the JSD (46), which is a symmetrized and smoothed version of the Kullback-Leibler divergence (46), given by,

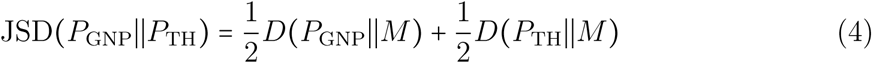

where *M* (*x, y*) = 1/2[*P*_GNP_(*x, y*) + *P*_TH_(*x, y*)], and *D* is the Kullback-Leibler measure given by,

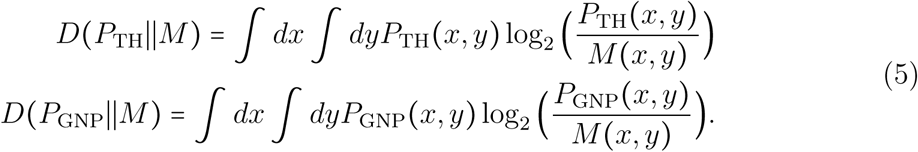

The value of the JSD measure varies between 0 and 1: the larger the value of the divergence, the greater is the difference between the two distributions. Both measures (K-S scores show a similar trend (see figure 2e) - as the size of the GNP increases, the difference between the distributions of the GNP and the TH increases.

### Attachment of the GNP increases the Rightward Bias

One of the most insightful findings of the experiments performed by Isojima *et al.* (35) is that the detached head undergoes a highly diffusive movement (see figure 2 in (35)). This observation, which was inferred by following the dynamics of the GNP, finds support in the simulations (31, 43). More importantly, it was noted by explicitly tracking the time-dependent trajectory of the GNP that the TH movement exhibited a rightward bias. This inference was based on the observation that the fluctuations transverse to the MT axis (see figure 1) are predominantly localized within the half-plane *Y*_*GNP*_ < 0.

In order to assess whether the GNP artificially introduces a rightward bias, we calculated the two-dimensional distribution *P* (*x, y*) = *P* (*X*_*TH*_ − *X*_*LH*_, *Y*_*TH*_ − *Y*_*LH*_) by averaging over the simulation trajectories. The plot in figure 3a, obtained in the absence of GNP and the cargo (figure 1) shows that the TH samples both the positive and negative values of *Y*_*TH*_ − *Y*_*LH*_. To quantify the results in figure 3, we calculated the asymmetry parameter,

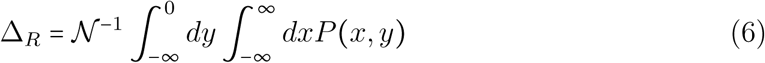

where 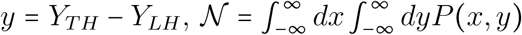. Note that Δ_*R*_ is bound between 0 and 1, where 0 (1) means that the tethered head overcomes the MT-bound motor always on the left (right). For the distribution in figure 3a, we found that Δ_*R*_ = 0.54, which implies during a step and in the absence of GNP, there is an almost equal probability that the TH is found on the right or on the left of the LH.

Attachment of GNP of various sizes enhances the rightward bias in the stepping trajectory, as illustrated in figures 3b and 3c. The value of Δ_*R*_ changes to 0.56 and 0.63 for the 10 nm and 20 nm GNP, respectively (see Table S1). Attaching a 40 nm GNP slows down the stepping process to such an extent that in our simulation time scale (≈ 20 *µ*s) NL docking of the LH is incomplete. As a consequence excursions of the GNP or the TH away from the initial binding site are small (see figure S7). From the increase in Δ_*R*_, as the GNP diameter increases, we surmise that the finding that TH always passes the LH on the right might be (at least partially) a consequence of attaching the GNP.

The 2D distributions, *P* (*X*_*GNP*_ − *X*_*LH*_, *Y*_*GNP*_ − *Y*_*LH*_), probing the movement of the 10 nm and 20 nm diameter GNPs, are shown in figure 3d - 3e. The comparison of the results in figure 3d and 3f shows that the rightward bias increases with the size of the GNP, leading once again to the conclusion that the GNP masks the true diffusional motion of the unbound head. Interestingly, the Δ_*R*_ increase is smallest when the GNP is attached to residue 55 (see figure 3f) but changes substantially for other attachment points (Table S1).

### How do changes in the GNP attachment positions affect the stepping trajectories

Isojima *et al*. (35) probed the movement of the free kinesin head by attaching the GNP to residues 55, 215, and 356, whereas Mickolajczyk *et al.* (34) adopted an N-terminal (residue 1) tag to attach the probe. We investigated whether these geometries enhance the rightward biasing by simultaneously tracking the position of the 10 nm GNP and TH relative to the bound head. Both the K-S score and the JSD show that the distributions of TH and GNP differ, and that the discrepancy increases with the size of the GNP (see figure 4a,b). This is most clearly shown by the JSD (figure 4a); the K-S score in the *X* directions increases significantly with the GNP diameter for all attachment sites, while the direction of the changes in K-S score with the size of the probe differ for the alternative attaching sites.

In all cases but one, the increase of the GNP size correlates with an increase of the TH rightward bias (see figure 4c). In addition, the behavior of the GNP depends on attachment site of the probe. These two observations underscore the difficulties highlighted by our study: if different attachment sites lead to different results (see figures 4-5, and figures S1-S2) and alter the trajectory of the TH in a system-dependent way (figure 4), it becomes extremely challenging to infer the dynamics of the TH using these massive probes. The results here show that the attachment point for the GNP has to be chosen judiciously to minimize biases in the stepping process.

**Figure 5:**
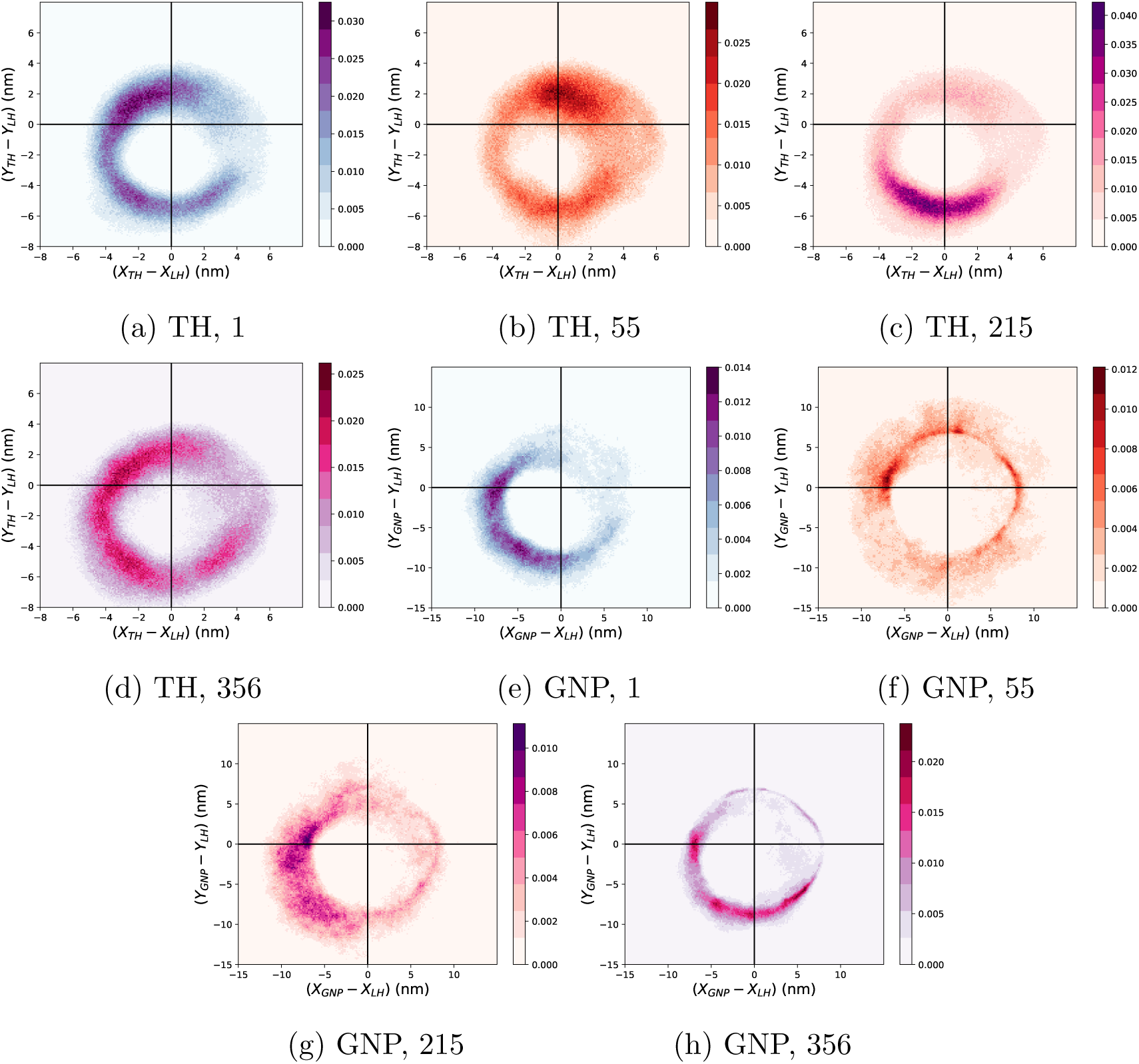
Two dimensional distribution of the position of the TH and GNP relative to the bound head. (a-d) Two-dimensional distribution of the position of the TH of the motor relative to the bound head with GNP (of size 10 nm) attached at residue 1 (a), 55 (b), 215 (c), and 356 (d). (e-h) The density plot of the position of GNP (size 10 nm), relative to the bound head, when attached at residue 1 (e), 55 (f), 215 (g), and 356 (h).

### Effect of cargo on the stepping of the TH and the GNP

In typical optical trap experiments a ≈ 1*µ*m bead is attached to the coiled coil. The movement of the cargo as kinesin steps on the MT is used as a proxy for the stepping kinetics. Although it could be difficult to realize experimentally, we assembled a construct with both a 1 *µ*m cargo attached to the coiled-coil and a GNP attached to residue 55 of the TH (figure S3).

We compared the resulting trajectories with those obtained in the presence of cargo but without the GNP. First, we observe that the rightward biasing of the TH trajectory is much more prominent in the presence of cargo (Δ_*R*_ = 0.83, see figure 6a) compared to the system without cargo (Δ_*R*_ = 0.55, see figure 3a). Interestingly, attaching a GNP to a cargo-bound motor reduces the rightward bias (Δ_*R*_ = 0.78 for 10 nm-GNP, and Δ_*R*_ = 0.74 for a 20 nm-GNP, see figure 6f). Second, when cargo is attached to the system, the NL of the LH remains docked throughout the stepping process. In other words docking is irreversible (figure S5a). In contrast, without cargo the NL docking of the LH is reversible in the time scale of our simulations (see figure S5b). Third, in the presence of cargo the mean docking time increases only slightly when the GNP is attached (≈ 8% for 10 nm GNP, and 4% for 20 nm GNP, see Table S2), whereas it grows significantly in the absence of cargo (≈ 60% for 10 nm GNP and 64% for 20 nm GNP).

**Figure 6:**
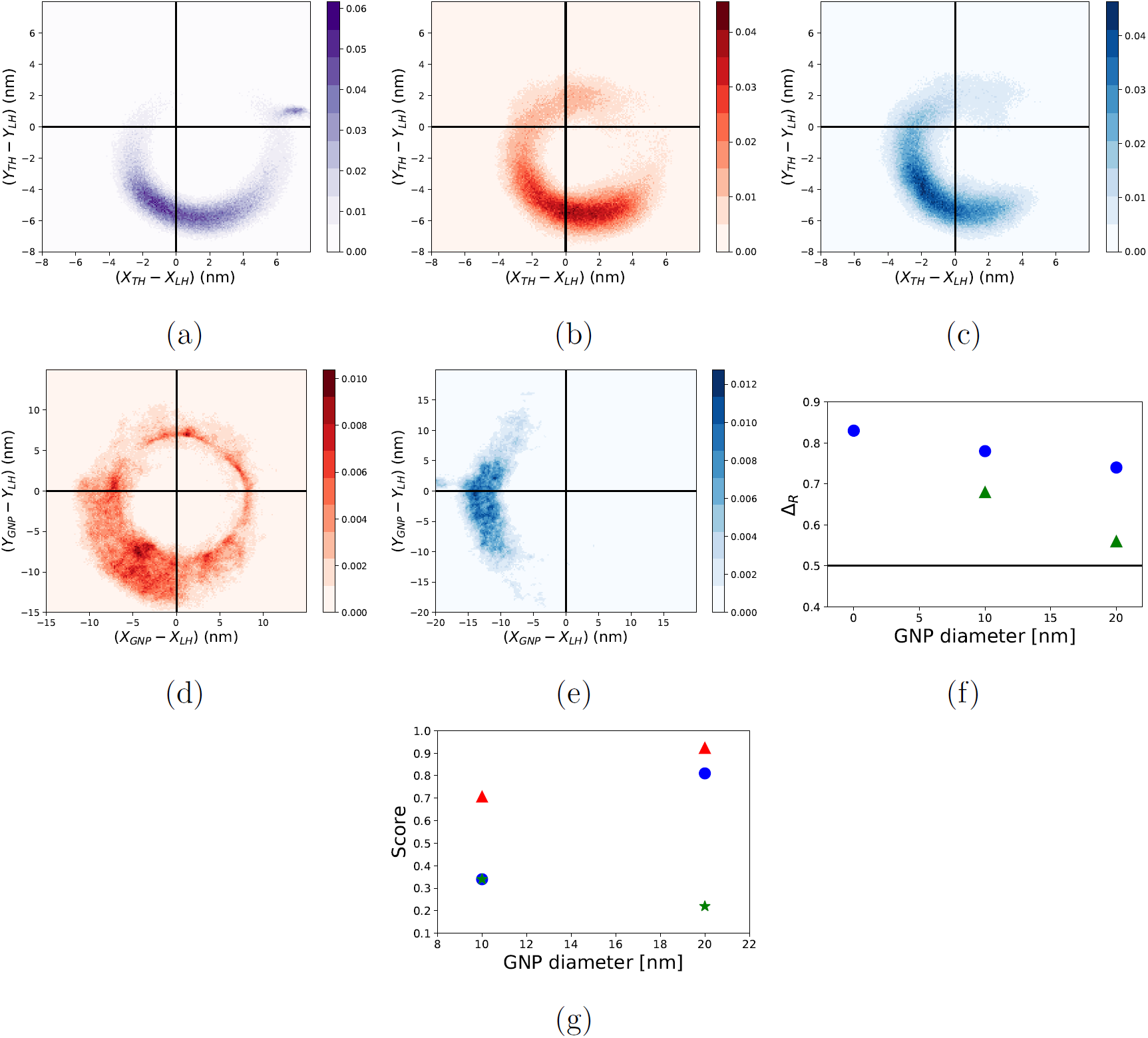
Comparison of the TH and GNP trajectories in the presence of cargo. (a-c)Two-dimensional density plot of the position of the TH with respect to the bound head for the system with cargo attached to the end of the coiled-coil (see figure S3 in the SI for the setup). The black lines in each figure correspond to the X and Y-axis with bound head located at position (0,0). The scale for the frequency of the distribution is shown in the right side of each plot. Distribution of the position of the TH of the motor with respect to the bound head for systems without GNP (a), 10 nm GNP (b), and 20 nm GNP (c). Distribution of the position of the GNP with respect to the bound head for systems with (d) 10 nm GNP (e) 20 nm. (f) Δ_*R*_ values for the TH (blue circles) and GNP (green triangle) for the system with cargo attached to the MT-kinesin complex. The black line shows Δ_*R*_ = 0.5.(g) Scores of the KS test and JSD when comparing the distribution of the positions of TH and GNP for the system with cargo. The circles in blue show KS test for the distribution of the positions along the X-direction. The star in green are the KS test scores for the distribution along Y-direction. Red triangle shows the JSD score (for the system with cargo).

Finally, we computed the JSD and K-S scores (discussed previously) for the system in the presence of both cargo and GNP (figure 6(g)). Even in this case, larger GNPs provide a poorer characterization of the position of the TH (see figure 6g), although the K-S score performance in the *Y* direction improves with the size of the GNP, as previously observed as well (see figure 2c and 2d).

## Conclusions

Motivated by the high-resolution experimental results obtained using iSCAT (34) and DFM measurements (35), we investigated how attaching a GNP to different residues on the tethered head of a kinesin motor impacts the stepping trajectories. This assessment cannot be conducted using experiments alone because it requires monitoring the positions of both the GNP and the motor head. Inferring the stepping dynamics of the motor from iSCAT or DFM experiments hinges upon the assumption that the GNP probe is an adequate proxy for the stepping head. That the GNP could alter the stepping trajectories can be anticipated by noting that the volume of the ≈ 30 nm GNP is roughly 1,000 times the volume of the motor head. From our simulations, which allow us to monitor the dynamics of both GNP and the TH *simultaneously*, we draw a few important conclusions that may be relevant to experiments on kinesin as well myosin V whose stepping has also been probed using GNP tracking (42, 47).

1. The interpretation of the experimental results is based on the assumption that the GNP does not significantly alter the stepping behavior of the motor. This observation is generally adduced by noting that the mean velocities of kinesin as a function of ATP concentration ([ATP]) for GNP and the fluorescently labeled motors do not differ significantly (35). For instance, the supplementary figure 1 in (35) shows that, in the absence of a resistive force, there is only ≈ 25% difference in the mean velocity measured using GNP-labeled motor head and flourescently labeled motor head at near physiological ATP concentration. The *V*_*max*_ value obtained by fitting the [ATP]-dependent velocity to Michelis-Menten equation differs by about 80 nm/s (*V*_max_ ≈ 615 nm/s for the fluorescently-labeled kinesin and is about 695 nm/s for the motor head labeled with GNP (see figure S1 in (35)). However, even if this difference is deemed to be small, it should be appreciated that the average motor velocity is likely to be insensitive to the microscopic details of the motor movement. This is because the GNP used by Mickolajczyk *et al.* (34) and Isojima *et al.* (35) are not large enough to change (or slow-down) the rate limiting step of kinesin at high ATP concentration. In order to understand why, let *V*_max_ ≈ 1 *µ*m/s should be the maximum velocity of kinesin (a slight overestimate), with an average run length of *λ* ≈ 1 *µ*m with an average attachment time of *τ* ≈ 1 s (*V* ≈ *λ*/*τ*). A probe attached to a kinesin head would impact the velocity of the motor if 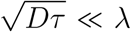, where *D = k*_*B*_ *T /(*6*πηR*) ≈ 0.25 (*µm*^3^ /s)*R*^−1^ is the diffusion coefficient of a sphere of radius *R* in water at room temperature (the viscosity of water is *η* ≈ 8.9 · 10^−4^ Pa·s, and *T* = 300 K). Therefore, only if the radius of the probe is ⪆ 0.25 *µ*m we expect to observe a significant reduction of the velocity due to the GNP. This is not the case in the current experimental setup used in iSCAT (34) and DFM (35). However, we do expect that the giant probe will modify the stepping mechanism of the motor. The reason is the following: the motility of a kinesin head is characterized by long dwells and fast displacements of Δ ≈ 16 nm, which may take a time as short as *τ* ≈ 20 *µ*s, as estimated from theoretical (48) and computational (31) studies. A GNP of *R* ≈ 20 nm is expected to traverse a distance of ≈ 16 nm in a time *τ* ≈ 20 *µ*s, and it therefore may be large enough to alter the fast step of kinesin. Furthermore, it is possible that the probe will interact with the motor tail, the bound head, or the MT. These interactions and the slow diffusion of the GNP could impact the dynamics of the motor without affecting the rate-limiting process, thereby making the interference of the GNP impossible to observe if only the maximal velocity is monitored. In accord with these arguments, our simulations show that the presence of the GNP alters the dynamics (e.g. increases the mean docking time) and trajectory of the tethered head.
2. An important conclusion in the experiments (35) is that the unbound head undergoes diffusive motion with substantial rightward bias. This implies that the transverse fluctuations should have predominantly negative *Y*_*TH*_ values in the coordinate frame shown in figure 1. The quantitative measure used in Eq. 6 shows that the probability of finding the TH with negative *Y*_*TH*_ does increase as the diameter of the GNP increases. The extent of increase in Δ_*R*_ depends on the residue location to which the GNP is attached. For the 20 nm GNP attached to resdue 55 there is nearly a 15% increase in the rightward bias when compared to the case with no GNP (Table S1). If the GNP is attached to residue 356 then there is 30% increase (Table S1). We surmise that the Δ_*R*_ value for commonly used the 40 nm GNP would increase even further. Because the simulation times for this case are too long we could not provide accurate value for Δ_*R*_.
3. Surprisingly our simulations reveal that if cargo is attached to the coiled coil in addition to the GNP (see figure S2 for the MT-kinesin system with cargo and GNP attached) then the mean docking time, the excursion of the TH along the MT axis at the time of docking, as well as transverse fluctuations do not change significantly (see Table S3 in the SI). In addition, the unbound head does pass the LH on the right with the values of Δ_*R*_ (Table S4 in the SI) that are similar with and without GNP. It is likely that the larger size of the cargo (diameter 1 *µ*m) compared to the sizes of the GNP implies that the TH movement is controlled by the cargo dynamics. It would be interesting to test our predictions by constructing a system with cargo and GNP attached to the coiled-coil and the TH, respectively. However, there could be technical difficulties in performing such experiments and in analysis of the stepping trajectories because of large hydrodynamic flows around the cargo and the GNP.

Recently, Mickolajczyk *et al*. (49) addressed the differences between the two interpretations of the nature of the ATP-waiting state previously proposed (34, 35), which is discussed in the Introduction. Using a combination of simulations and experiments, the authors conclude that the discrepancies are a byproduct of the different labeling geometries used by the two groups. Importantly, they reiterate that kinesin steps on the right of the bound head, and they propose that the motor awaits for ATP binding in a conformation denominated as “unbound-undisplaced,” in which the tethered head is detached from the MT but “hovers” in the vicinity of the initial binding site. In comparing with our work, it is first necessary to notice that our model is much more descriptive, as it includes structural details of the motor, the NL docking, MT etc. that are not accounted for by Mickolajczyk *et al*. (49). Secondly, Mickolajczyk *et al*. (49) construct their model in order to reproduce the observed time-traces of previous experiments, including the rightward bias of the stepping head, and therefore do not address the questions that motivated our study. In particular, they do not systematically examine the bias introduced by the GNP and whether the GNP faithfully tracks the location of the TH. Therefore, our investigation is complementary to the work of Mickolajczyk *et al*. (49). Thirdly, in the original study of the kinesin stepping model adopted here (43), a mutant incapable of docking the NL was considered. The simulations of this mutant provided a representation of the conformations explored by a tethered head in the 1HB, ATP-waiting state (ATP binding induces NL docking). The resulting ensemble of structures qualitatively differs from the “unbound-undisplaced” picture proposed by Mickolajczyk *et al*. (49), because the tethered head was found to move away from the original binding site. Therefore, the physical basis of the interactions required to stabilize the “unbound-undisplaced” state need to be determined. Finally, a recent study employing a kinetic model of kinesin showed that the randomness parameter could be used in order to distinguish between the 1HB and 2HB models for the ATP-waiting state (50). The advantage of this approach is that it involves a measurement which does not require the use a GNP as a probe, and therefore constitute a test independent of the insights drawn from iSCAT and DFM studies.

## Model and Methods

### Self-Organized Polymer (SOP) model

We used the SOP model for the MT-kin complex (44, 51–53), which was introduced elsewhere (31, 43) in order to characterize the stepping dynamics of conventional kinesin. Unlike atomically detailed simulations (29), our studies, based on the coarse-grained SOP model, reproduce accurately experimental time-dependent changes in the position of cargo subject to resistive force of 5 pN (see figure 1 in ref. (22)), thus providing a measure of validation. In the SOP model, each residue in the MT-Kin complex is represented by single bead of diameter 0.38 nm that is located at the *C*_*α*_ atom. As in the previous studies (31, 43), we included three MT protofilaments (see figure 1), two motor heads, a coiled coil whose contour length is 30 nm.

The SOP energy function used to describe the MT-kin complex is given by,

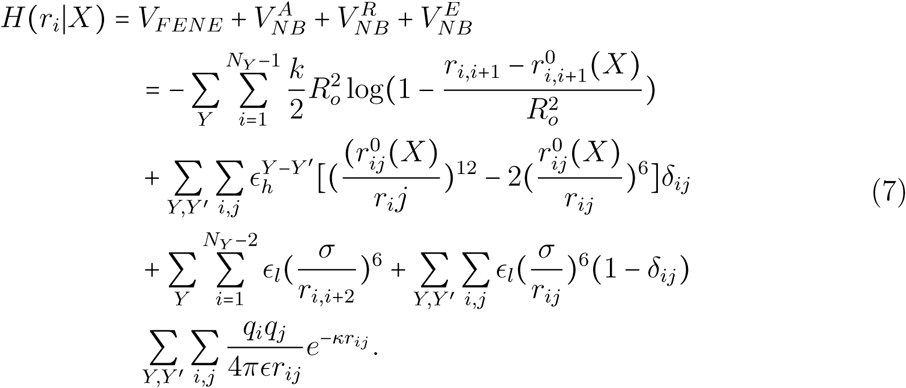

The first term in Eq. 7 describes the intramolecular connectivity of the molecule, which is given by the finite extensible non-linear elastic potential (FENE). The second term accounts for attractive interactions between the residues that are in contact in the type, low-energy MT-Kin complex. The third term corresponds to the repulsive interactions between residue pairs that are not in contact in the MT-kin complex, and the fourth term corresponds to the electrostatic interaction between pairs of charged residues. The symbols *Y* and *Y′* refer to the residues in the leading head (LH), trailing head (TH) and the MT. The argument *X* represents the state describing different stages of the two motor heads. One is the two head bound ATP-waiting state in which the NL is docked to TH while the NL is undocked in the LH. The other is the stepping state in which the NL docked to the LH after ATP binding, and the NL undocks from the TH. The values of the interaction strengths determining the NL docking to the LH and the TH interaction with MT were taken from our previous study (31).

In the SOP model there are two energy scales that determine the stepping kinetics of kinesin, which were calculated in order to obtain accurate values of the stall force (*ϵ*^*LH* − *NL*^) and the force required to unbind a single motor head from the MT (*ϵ*^*MT* − *TH*^). The value of *ϵ*^*LH*^ − ^*NL*^ is taken as 0.3 kcal/mol, and the value of *ϵ*^*MT − TH*^ is 0.2 kcal/mol (31). At the start of the stepping process (t=0) interaction (*ϵ*^*LH* − *NL*^) between the residues of the NL of the LH (326-338) and the residues of the LH are taken to be attractive provided the the criteria |*i* − *j*| > 2 and 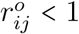 nm is satisfied. This ensures that the NL docks to the LH. The binding of the TH to the MT at the completion of the stepping is favored by the attractive potential, with strength *ϵ*^*MT − TH*^ between the TH and the binding site on the MT. The attractive interactions for the NL docking of the LH and binding of the TH to the MT is calculated using the second term in Eq. 7. Additional details of the SOP energy function may be found elsewhere (31, 43)).

### Modeling the GNP and the tether between the TH and the GNP

We performed simulations by attaching the GNP through a tether (a polymer chain) to a residue Let us first describe how the GNP is attached to a specific residue on the motor head. through the tether (figure 1). The polymer chain has 17 monomers, mimicking the Maleimide-PEG2-biotin (size ≈ 2.9 nm), and roughly the size of the biotin-streptavidin complex (≈ 3.35 nm). One end of the polymer (bead 1) is attached to the GNP, and the other end (bead 17) is attached to one of the residues (1, 55, 215, or 356) in the motor head. The attachment of the first bead on the polymer to the GNP is governed by a harmonic potential,

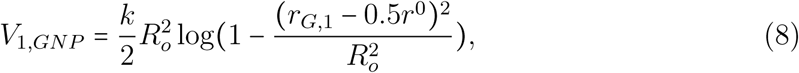

where *R*_*G*_ is the radius of the GNP, and *r*_*G*,1_ is the distance between the center of the first bead to the surface of the GNP. Similarly, a harmonic potential is used to attach bead 17 on the polymer to the residue on the motor head. We set the diameter of each bead, *r*^0^ = 0.38 nm, which implies that the contour length of the polymer is 6.08 nm (= 16 × 0.38 nm). The value of the spring constant for polymer is taken by FENE potential, *k =* 2, 000 kcal/(mol ^2^). The FENE potential (first term in Eq. 7) is used to ensure connectivity of the polymer beads 2 through 16.

The non-bonded interactions between the polymer beads are modeled using a purely repulsive interaction whose functional form is given by the third term on the right hand side of Eq. 7. The repulsive interactions of the beads in the polymer with the residues in the TH, LH, MT, and charged E-hooks in the MT are given by,

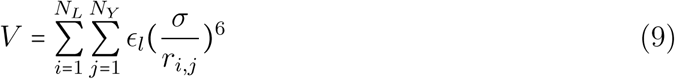

where Y corresponds to the residues in TH, LH, MT, E-hooks; *N*_*L*_ is the number of monomers in the polymer, *N*_Y_ is the number of the residues in *Y, σ =* 0.38 nm and *ϵ*_*l*_ = 1 kcal/mol. The volume exclusion between the GNP with every other particle is given by repulsive LJ potential described in Eq. 10 with *σ =* 0.38 nm from the surface of the GNP.

The repulsive interaction between the GNP and the interaction sites on the TH, LH, polymer, MT, and E-hooks is given by,

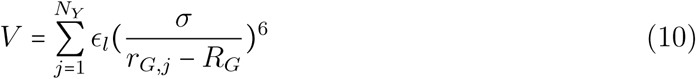

where Y labels the residues in TH, LH, the polymer, MT, E-hooks and *N*_*Y*_ corresponds to the number of residues in Y. In Eq. 10 *r*_*G,j*_ is the distance between the residue in Y from the center of the GNP.

### Simulations

In order to elucidate the consequences of attaching the GNP of differing sizes (diameter ranging from 10 − 40 nm) to various residues on the TH, we performed a number of simulations. Except for the treatment of the GNP, which we describe in detail below, we followed the procedure described in our previous studies (31, 43). We assume that the stepping trajectories could be generated by integrating the Langevin equations in the over-damped limit for the MT-kin complex both in the absence and presence of the GNP.

Hydrodynamic interactions were included for residues in TH, polymer, and the GNP using the Rotne-Prager-Yamakawa tensor approximation (54). We also took into consideration the rotation of the spherical GNP while it undergoes translational motion (see below).

### GNP dynamics

Because the GNP is tethered to a polymer, which in turn is linked to specific residues on the TH, its dynamics has to be treated with care. The GNP is spherical and in isolation would undergo isotropic rotation. However, by attaching the GNP through the tether the motion of the composite systems (GNP, tether, and the motor) is inherently anisotropic. Consequently, the GNP undergoes both translational and a rotational motion while it is covalently attached to the one end of the polymer with the other end connected to the TH of the motor (Eq. 7). We treat the translational and rotational motion independently. At the start of the simulations (t=0), the polymer bead labeled 1 is attached to a point on the surface of the GNP. Because of the spherical symmetry, the initial orientation vector (k) at time *t* = 0 can be in any direction. In our simulations we chose k=(1,0,0) at *t* = 0.The objective is to obtain the position of the attachment (see figure 1) as a function of time.

The rotation of the GNP, with respect to the tether and the TH, changes the orientation of the sphere, which is accommodated by taking a point particle at the surface of the GNP, and tracking its angular displacement. We calculated the vector *r* from the center of the GNP to the surface (point of attachment of the tether to the surface of GNP) (figure 7). The torque at time *t* is calculated using, *T = r × F*, where *F* is the force acting on the surface of GNP in contact with the polymer bead. The orientation (*k* (*t* + Δ*t*)) after a time Δ*t* (integration time step in the equation of motion) is elapsed is given by, (55)

**Figure 7:**
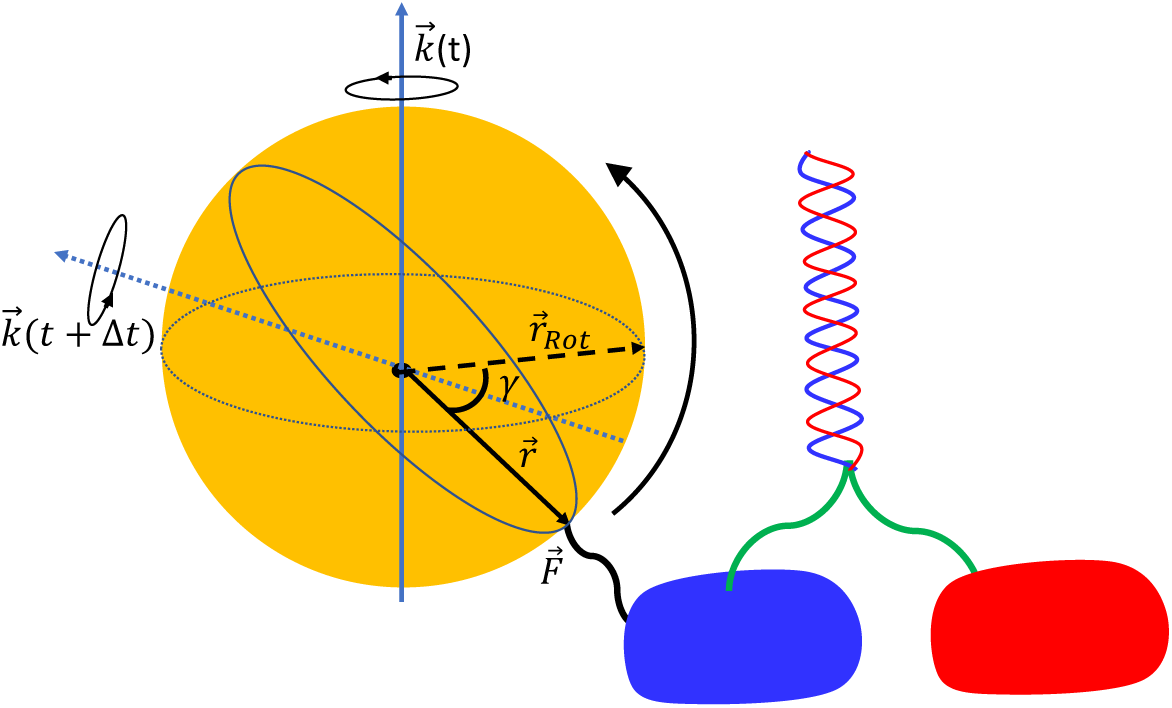
Change in the orientation of GNP during rotation. Here, *k* (*t*) and *k* (*t* + Δ*t*) are the rotation vectors at time *t* and *t* + Δ*t, γ* is the rotation angle, 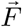 is the force at the attachment point of GNP surface to the polymer bead, 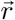 is the vector from the center of the GNP to the surface and 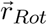 is the vector obtained after rotation.

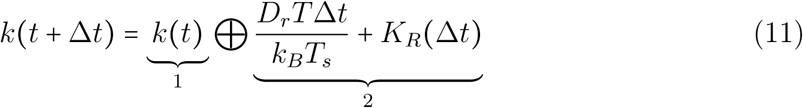

where *k* (*t* + Δ*t*) is the orientation at time *t* + Δ*t, k* (*t*) is the orientation at time *t, T* is the value of torque at time *t, D*_*r*_ is the rotational diffusion coefficient of GNP 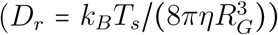, *η* is the viscosity, *k*_*B*_ is the Boltzmann constant, *T*_*s*_ is the simulation temperature, and *K*_*R*_ (Δ*t*) is a random rotational step. We sampled *K*_*R*_ from a Gaussian distribution with zero mean and standard deviation (2*D*_*r*_Δ*t*)^1 2^. The sum of the last two terms in Eq. 11 is a vector sum, whose length is the rotation angle (*γ*) and the direction is the rotation axis. This vector is converted to rotation vector, with magnitude *tan* (*γ /*2). The ⊕ is a symbol for adding two rotation vectors, and is given by (55)

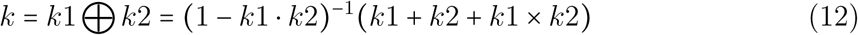

where *k* is the resultant rotation vector, × is the cross product between the two vectors. The resultant rotation vector obtained from Eq. 12 is transformed to rotation matrix, the elements of which are given by,

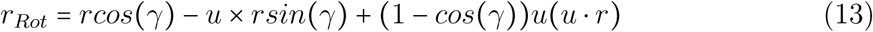

where *r*_*Rot*_ is the vector after transformation, and *r* is the initial vector, and *u* = *k* /|*k*|. The position of the attachment of the polymer to the surface of GNP after rotation is updated is given as,

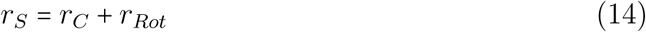

where *r*_*S*_ is the coordinate of the point of attachment of the polymer on the surface of GNP, and *r*_*C*_ is the position of center of GNP.

### Simulations with cargo attached to the motor

We also performed simulations to assess the effect of cargo on the MT-kin system in the presence of GNP. The cargo, diameter 1 *µ*m, is attached to the end of the coiled-coil, as in our previous studies (31, 43). The cargo is connected by the FENE potential (given by the first term in Eq. 7) to the end of the coiled-coil, and volume exclusion is taken using repulsive LJ-potential (given by Eq. 10) with the motor, GNP, and the polymer tag. **Simulation Details:** In generating every stepping trajectory we first equilibrated the system of interest (for example the MT-Kin1 complex with GNP attached to a specific residue) for 500,000 time steps, which corresponds to 0.5 *µ*m. On this time scale the GNP has moved roughly 10 nm. Thus, there is no memory of initial conditions when we generate the stepping trajectory. Further details are in the SI.

## Supporting Material

Supporting material is provided.

## Acknowledgments

This work was supported in part by a grant from National Science Foundation (CHE 19-00093) and the Collie-Welch Chair (F-0019) administered through the Welch Foundation.

