## Supplementary material for "Effects of Gold Nanoparticles on the Stepping Trajectories of Kinesin": Sup

February 2, 2020

**Additional simulation details:** For each set up, we first performed the equilibrium simulation for $5 \times 10^6$ time steps, which is equivalent to $0.5\,\mu$s. During the equilibrium simulations, the motor remains in two head bound state with TH in the docked state and LH in undocked state. The TH of the motor is bound to $\alpha\beta$ tubulin at the (-) end of the central protofilament and LH to the middle of the $\alpha\beta$ tubulin of the same protofilament.

After equilibration is complete, the motor switches from the 2H bound state to 1H state in which LH is strongly bound to the MT. The stepping starts with the detachment of the TH of the motor from the MT because the repulsive interaction between the TH and the (-) end of the $\alpha\beta$-tubulin favours the detachment of the TH. The docking of the NL of LH is achieved by attractive interactions, $\epsilon^{LH-NL}$(0.3 kcal/(mol)), between the residues of the LH and the NL (T326-338). NL docking to the LH propels the TH towards the (+) end of the MT cite (**?** ).

**Order parameter for docking of Neck Linker (NL) of the Leading Head (LH):** The conformational changes that occur in the LH are used to monitor NL docking. We calculated the order parameter ($\Delta_{NL}\ (t)$) to assess if the NL is docked to the LH (S1),

$$\Delta_{NL}(t) = \sqrt{\left(\frac{\sum_{i,j}(r_{i,j}(t) - r^0_{ij})^2}{N}\right)} \tag{S1}$$

where $i$ and $j$ are the residue pairs involved in docking, $r_{ij}$ is the distance between the residues at time $t$, and $r^0_{ij}$ is the distance between them when the LH is in the docked state. The value of $r^0_{ij}$ is calculated using rat monomeric kinesin structure (PDB code 2kin). The summation in the above expression is over all $r_{ij}$ whose distance is less than 1 nm in the docked state. The NL of LH is assumed to be docked if $\Delta_{NL}(t) < 0.365$ nm. The NL docking times are calculated using the distribution of first passage times, which are obtained by generating a number of stepping trajectories. The first passage time for a given trajectory is identified with the first time a given trajectory satisfies the condition $\Delta_{NL}(t) < 0.365$ nm.

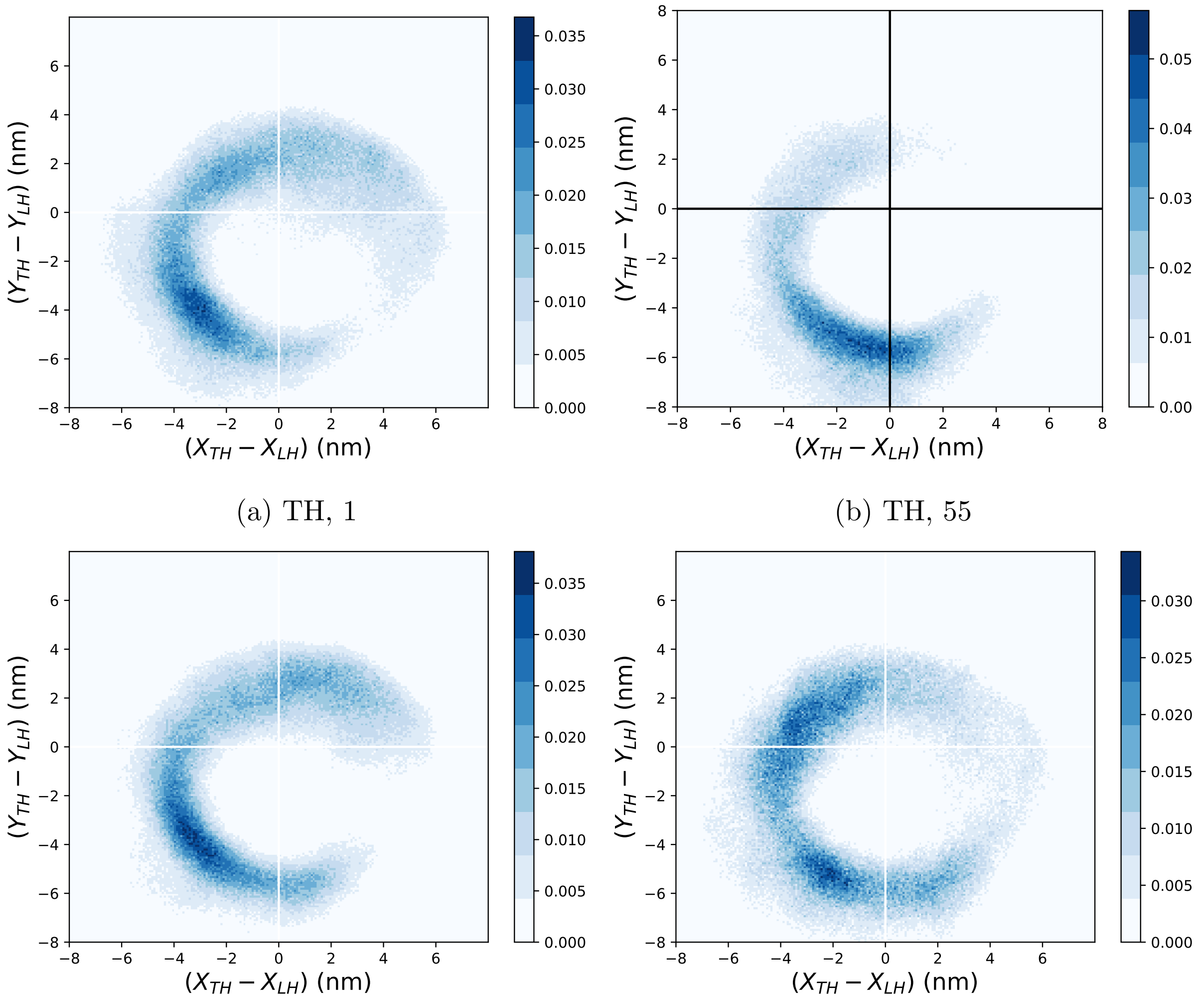


(a) TH, 1 (b) TH, 55

(c) TH, 215 (d) TH, 356

Figure S1: Two-dimensional distribution of the position of the TH of the motor (relative to the bound head) attached to GNP of size 20 nm. The center white lines indicate $X = 0$ and $Y = 0$. The scale for the intensity is given in the color bar on the right of each plot. Distribution of the position of TH relative to the (MT-bound) LH when 20 nm GNP is attached to residue (a) 1 (b) 55 (c) 215 and (d) 356.

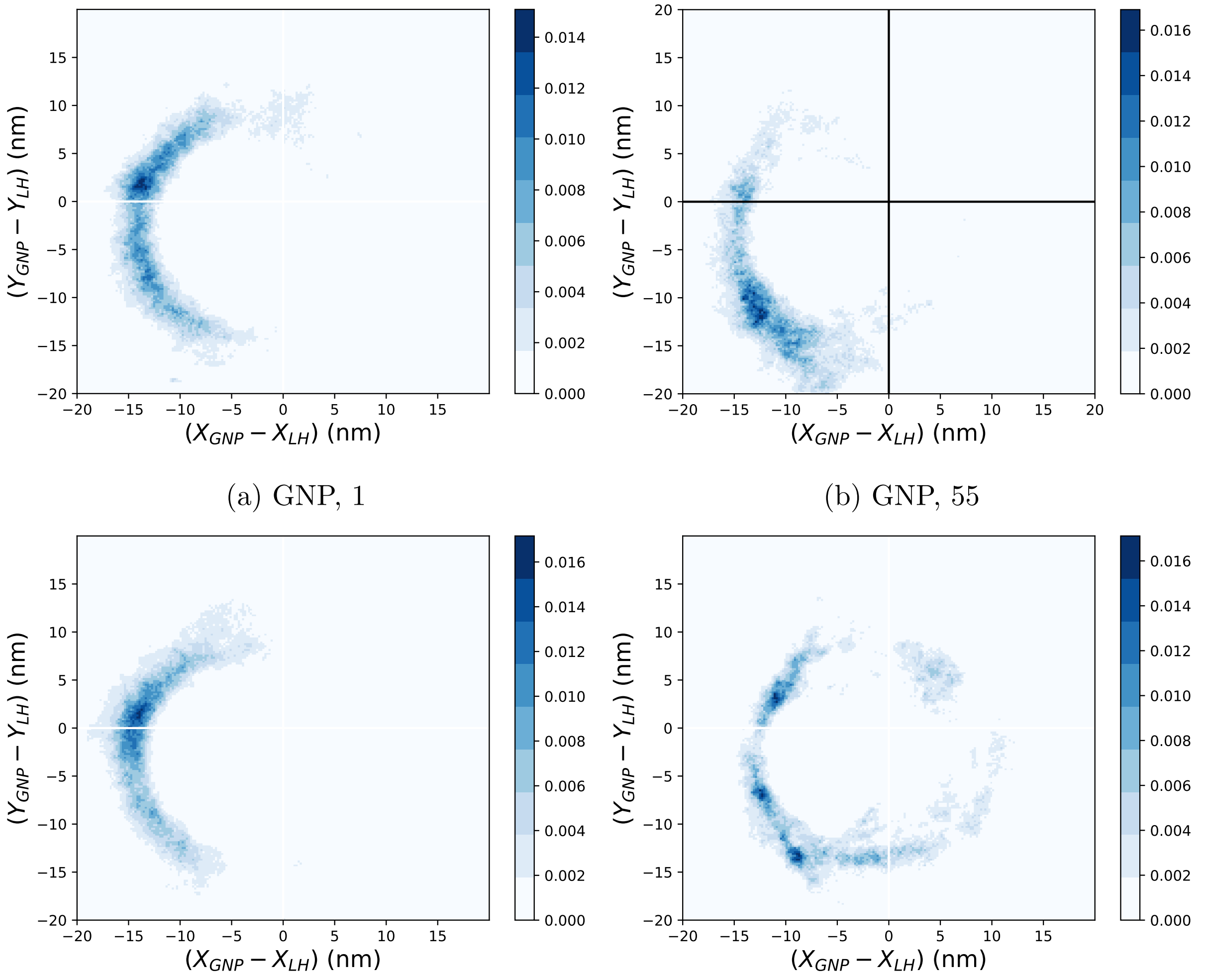


(a) GNP, 1

(b) GNP, 55

(c) GNP, 215

(d) GNP, 356

Figure S2: Two-dimensional distribution of the position of the GNP (20 nm) with respect to the LH. The center white lines indicate $X = 0$ and $Y = 0$. The scale for the intensity is given in the color bar on the right of each plot. Distribution of the position of GNP relative to the LH when attached to residue (e) 1 (f) 55 (g) 215 and (h) 356.

Table S1

| System | Attachment point | $\Delta_R{}^a$ | Standard Error (SE) |
|---|---|---|---|
| No GNP | - | 0.55 | 0.06 |
| 10 nm GNP | 1 | 0.57 | 0.07 |
| 10 nm GNP | 55 | 0.56 | 0.07 |
| 10 nm GNP | 215 | 0.73 | 0.06 |
| 10 nm GNP | 356 | 0.66 | 0.07 |
| 20 nm GNP | 1 | 0.60 | 0.06 |
| 20 nm GNP | 55 | 0.63 | 0.07 |
| 20 nm GNP | 215 | 0.59 | 0.07 |
| 20 nm GNP | 356 | 0.71 | 0.05 |
| 40 nm GNP | 55 | 0.44 | 0.07 |

a. The values are calculated using Eq. 6 in the main text for the set up that is illustrated in Fig. 1 in the main text.

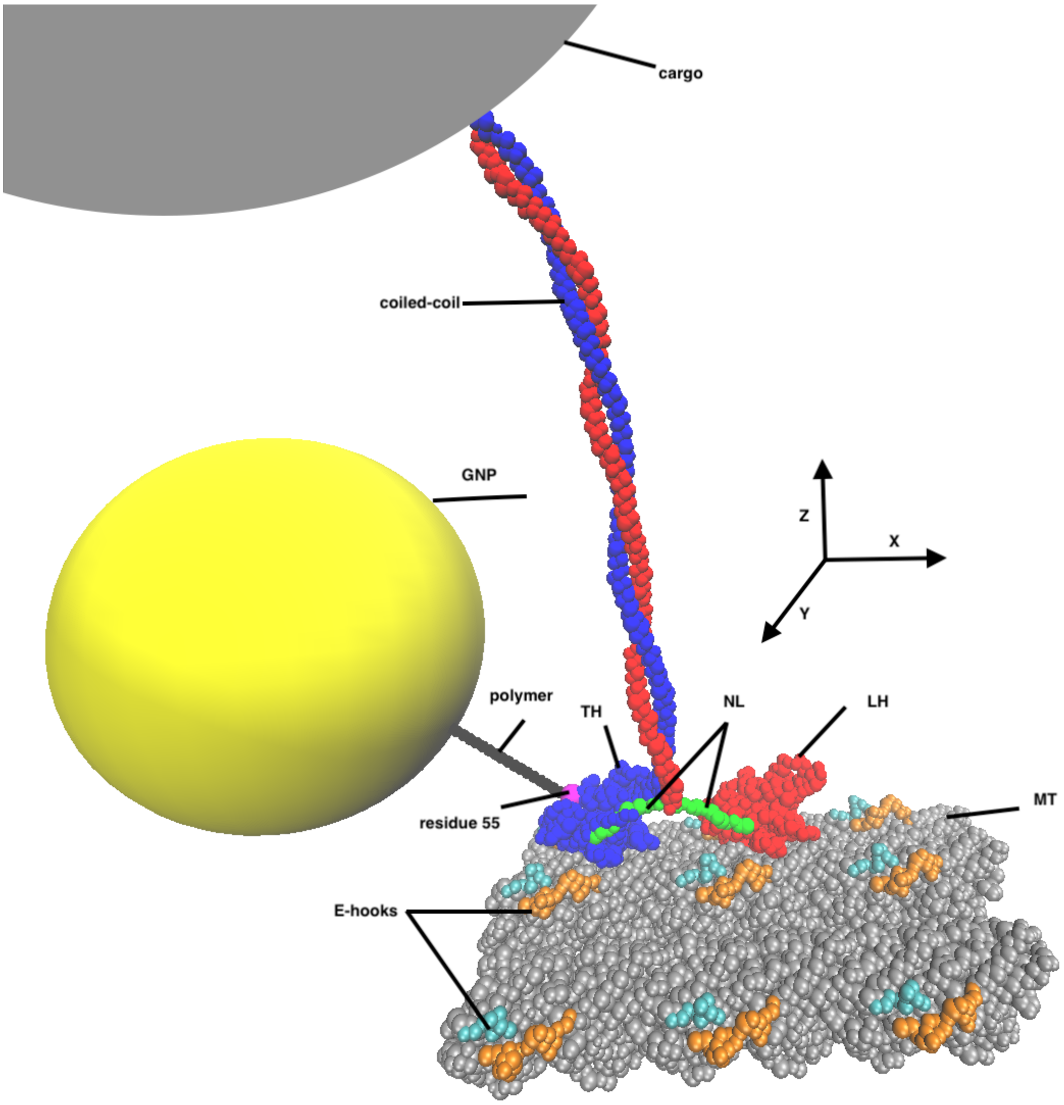


Figure S3: Schematic representation of the MT-kinesin complex with GNP (marked as yellow sphere) attached to the TH at residue 55 (marked in magenta) through a 17 bead polymer (marked in black). The cargo of diameter 1000 nm (shown as grey sphere at the tail) is attached to the end of the coiled-coil. The charged E-hooks are shown in cyan and magenta and MT is shown in silver color. The direction of stepping of the motor is along the X-axis marked and the lateral displacement correspond to the displacement along the Y-direction.

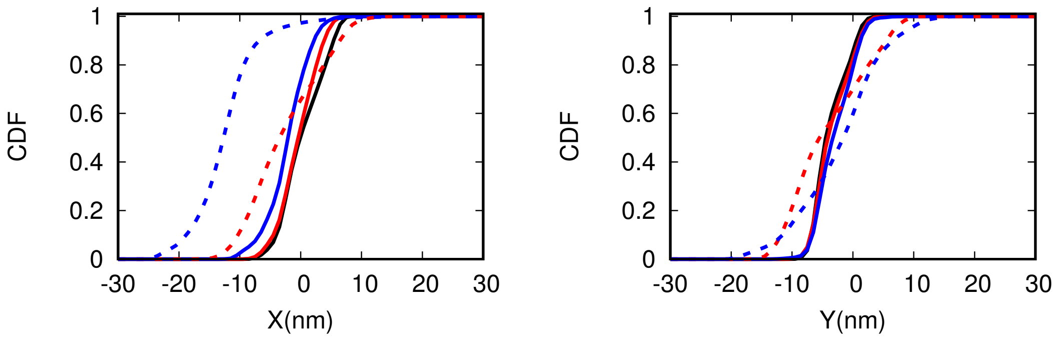


(a) X-direction (b) Y-direction

Figure S4: Cumulative distribution function (CDF) of the position of TH and GNP relative to the position of bound head along (a) X-direction and (b) Y-direction. The lines in dash is the CDF for the GNP, and the solid line correspond to the distribution of the TH with respect to the LH. The colors black, red, and blue correspond to the system without GNP, 10 nm GNP and 20 nm GNP system.

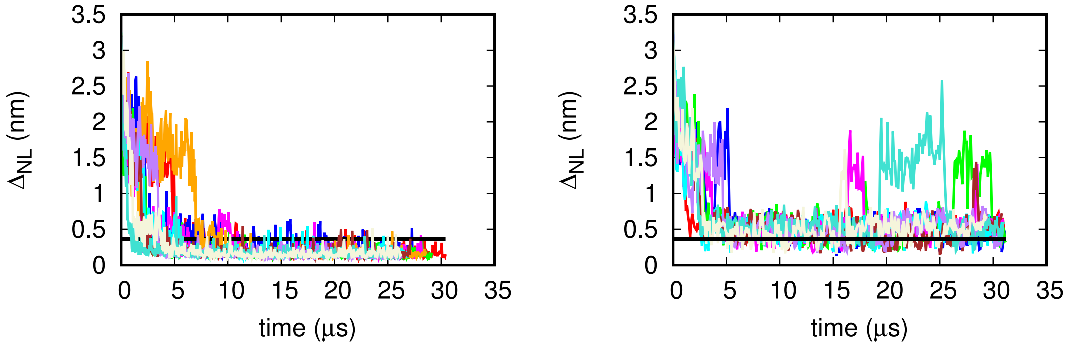


(a) with cargo (b) without cargo

Figure S5: Variation of the docking order parameter $\Delta_{NL}(t)$ (Eq. S1) with time for 10 trajectories for systems without GNP (a) with cargo, and (b) without cargo. The LH is docked if $\Delta_{NL}(t) < 0.365$. The black line in the figures corresponds to the $\Delta_{NL} = 0.365$. The figure on the right shows that NL fluctuates to the undocked state even after initial docking where as in the presence of cargo it is irreversible.

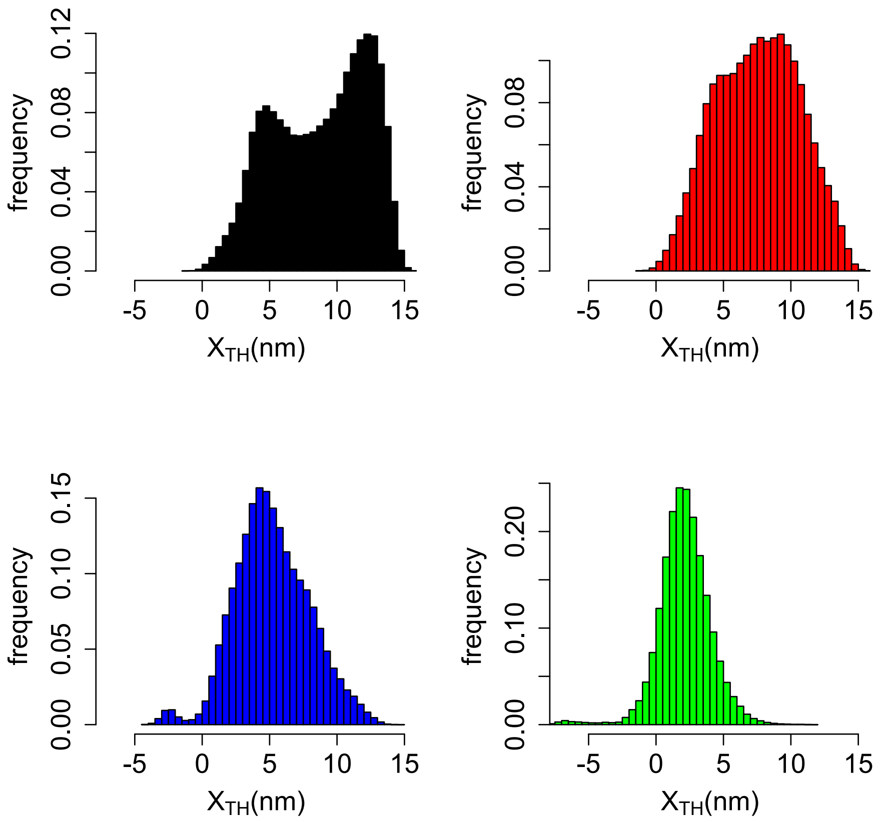


Figure S6: Distribution of the displacement of the TH ($X_{TH}$) along the MT-axis for the system without cargo. The figure in black, red, blue, and green correspond to the system without GNP, 10 nm GNP, 20 nm GNP, and 40 nm GNP, respectively.

Table S2: Mean NL docking time and the displacement of the TH at the instance of docking

| System | Attachment point | Time ($\mu$s) | $X_{TH}$ (nm) | $Y_{TH}$ (nm) | # of trajectories |
|---|---|---|---|---|---|
| No GNP | - | $3.87 \pm 2.45$ | $9.01 \pm 3.029$ | $-2.61 \pm 2.99$ | 69 |
| 10 nm GNP | 1 | $5.09 \pm 4.28$ | $7.18 \pm 2.76$ | $-1.15 \pm 2.82$ | 47 |
| 10 nm GNP | 55 | $6.33 \pm 4.50$ | $7.88 \pm 2.58$ | $-1.16 \pm 2.77$ | 48 |
| 10 nm GNP | 215 | $4.02 \pm 4.02$ | $3.79 \pm 3.04$ | $1.45 \pm 3.07$ | 49 |
| 10 nm GNP | 356 | $7.32 \pm 7.39$ | $4.76 \pm 2.96$ | $-0.44 \pm 2.92$ | 46 |
| 20 nm GNP | 1 | $6.79 \pm 5.14$ | $7.11 \pm 2.43$ | $-1.62 \pm 3.07$ | 43 |
| 20 nm GNP | 55 | $6.37 \pm 4.79$ | $5.77 \pm 1.87$ | $-1.68 \pm 2.72$ | 69 |
| 20 nm GNP | 215 | $7.49 \pm 6.10$ | $6.17 \pm 1.96$ | $-1.84 \pm 3.05$ | 49 |
| 20 nm GNP | 356 | $15.47 \pm 6.54$ | $8.47 \pm 2.81$ | $-1.66 \pm 4.02$ | 35 |
| 40 nm GNP | 55 | $11.40 \pm 7.14$ | $6.45 \pm 1.90$ | $-1.19 \pm 3.14$ | 17 |

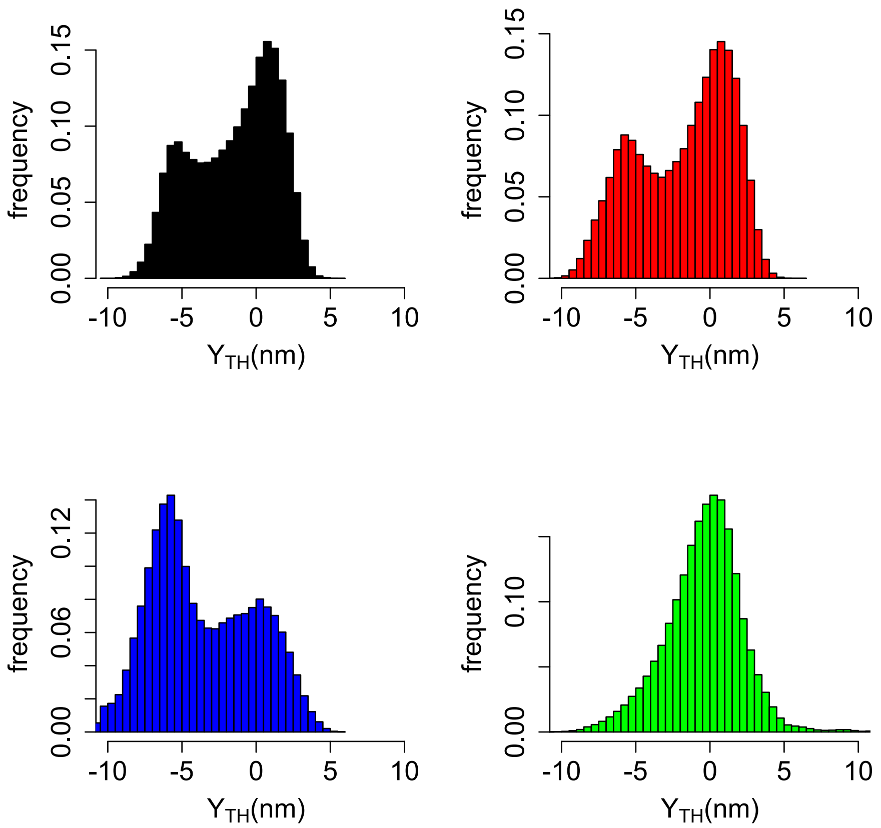


Figure S7: Distribution of the displacement of the TH ($Y_{TH}$) along the Y-axis for the system without cargo. The figure in black, red, blue, and green correspond to the system without GNP, 10 nm GNP, 20 nm GNP, and 40 nm GNP, respectively.

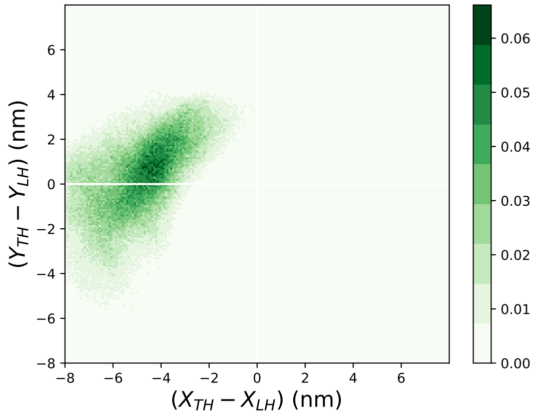


(a) TH

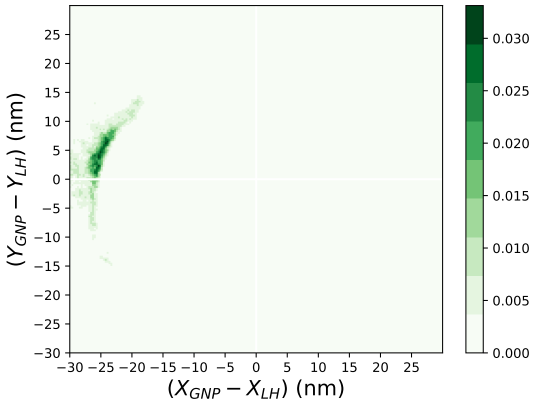


(b) GNP

Figure S8: Two dimensional density plot for the distribution of the position of (a) TH and (b) GNP with respect to the position of the bound head for the system with 40 nm GNP attached at residue 55 of the TH. The while lines in the figures correspond to X-and Y-axis. The bound head is located at the center (0,0) of the XY plane. The scale for the probability density is on the right.

Table S3: Mean docking time and the displacement of TH at the instance of docking (with 1000 nm cargo at the end of coiled coil (Figure S3).

| System | Attachment point | Time ($\mu s$) | $X_{TH}$ (nm) | $Y_{TH}$ (nm) | # of Trajectories |
|---|---|---|---|---|---|
| No GNP | - | $3.88 \pm 3.17$ | $7.39 \pm 2.83$ | $-3.88 \pm 2.80$ | 100 |
| 10 nm GNP | 55 | $4.19 \pm 2.85$ | $6.56 \pm 2.50$ | $-3.12 \pm 2.96$ | 50 |
| 20 nm GNP | 55 | $4.09 \pm 3.82$ | $5.17 \pm 1.71$ | $-3.36 \pm 2.61$ | 47 |

Table S4

| System | Attachment point | ${\Delta_R}^b(withcargo)$ | Standard Error (SE) |
|---|---|---|---|
| No GNP | - | 0.83 | 0.04 |
| 10 nm GNP | 55 | 0.78 | 0.06 |
| 20 nm GNP | 55 | 0.74 | 0.06 |

(b) The values are calculated using Eq. 6 in the main text for the set up that is illustrated in Figure S3.
